# Cell-type specific innervation of cortical pyramidal cells at their apical tufts

**DOI:** 10.1101/571695

**Authors:** Ali Karimi, Jan Odenthal, Florian Drawitsch, Kevin M. Boergens, Moritz Helmstaedter

## Abstract

We investigated the synaptic innervation of apical tufts of cortical pyramidal cells in a region between layers 1 and 2 using 3-D electron microscopy (3D-EM) applied to four cortical regions in mouse. Across all cortices, we found the relative inhibitory input at the apical dendrite’s main bifurcation to be more than 3-fold stronger for layer 2 pyramidal cells than for all other pyramidal cells. Towards the distal tuft dendrites in upper layer 1, however, the relative inhibitory input was about 2-fold stronger for L5 pyramidal cells than for all others. Only L3 pyramidal cells showed homogeneous inhibitory input density. The inhibitory to excitatory synaptic balance is thus specific for the types of pyramidal cells. Inhibitory axons preferentially innervated either layer 2 or L3/5 apical dendrites, but not both. These findings describe connectomic principles for the control of pyramidal cells at their apical dendrites in the upper layers of the cerebral cortex and point to differential computational properties of layer 2, layer 3 and layer 5 pyramidal cells in cortex.

## INTRODUCTION

The apical dendrites of pyramidal neurons, the most abundant cell type in the mammalian cerebral cortex (Cajal 1899, DeFelipe and Fariñas 1992), have been a key focus of anatomical (Cajal 1899), electrophysiological (Stuart and Sakmann 1994, Larkum, Zhu et al. 1999) and *in-vivo* functional studies (Svoboda, Denk et al. 1997, Letzkus, Wolff et al. 2011, Takahashi, Oertner et al. 2016) for their ability to electrically bind long-range top-down inputs converging at the apical tufts in layer 1 with synaptic inputs on the basal dendrites (Larkum, Zhu et al. 1999, Larkum 2013). Since almost all pyramidal cells from cortical layers 2, 3 and 5 (and some from layer 4) form their apical dendrite’s main bifurcation in this region, the layer 1/2 border in the upper cortex allows for synaptic convergence onto 80-90% of all cortical pyramidal cells (Larkman and Mason 1990, Ito, Kato et al. 1998). Furthermore, the apical dendrites’ main bifurcation has been implied as a site of origin and control of regenerative electrical events in the distal dendrites (Helmchen, Svoboda et al. 1999, Larkum and Zhu 2002, Larkum, Senn et al. 2004). Understanding the innervation profile of synaptic input onto apical dendrites in this peculiar part of the mammalian cerebral cortex was the goal of this study.

## RESULTS

We acquired four 3D-EM datasets from mouse somatosensory (S1), secondary visual (V2), posterior parietal (PPC) and anterior cingulate cortex (ACC) sized between 72 × 93 × 141 µm^3^ and 66 × 89 × 202 µm^3^ (Fig. 1a, Suppl. Fig. 1a,c, Suppl. Table 1) at a voxel size of 11.24-12 × 11.24-12 × 28-30 nm^3^ using serial block-face electron microscopy (SBEM (Denk and Horstmann 2004)). The image volumes were targeted at the border of layers 1 and 2, the site of the main bifurcation of apical dendrites from almost all pyramidal neurons of layers 2 to 5 (Fig. 1a,b). To determine the layer origin of each apical dendrite (AD), we first identified those ADs with a soma in the image volume as L2 ADs, and the remaining ADs as “deeper layer” (DL ADs), i.e. L3 and L5 ADs (Fig. 1a).

**Figure 1.**
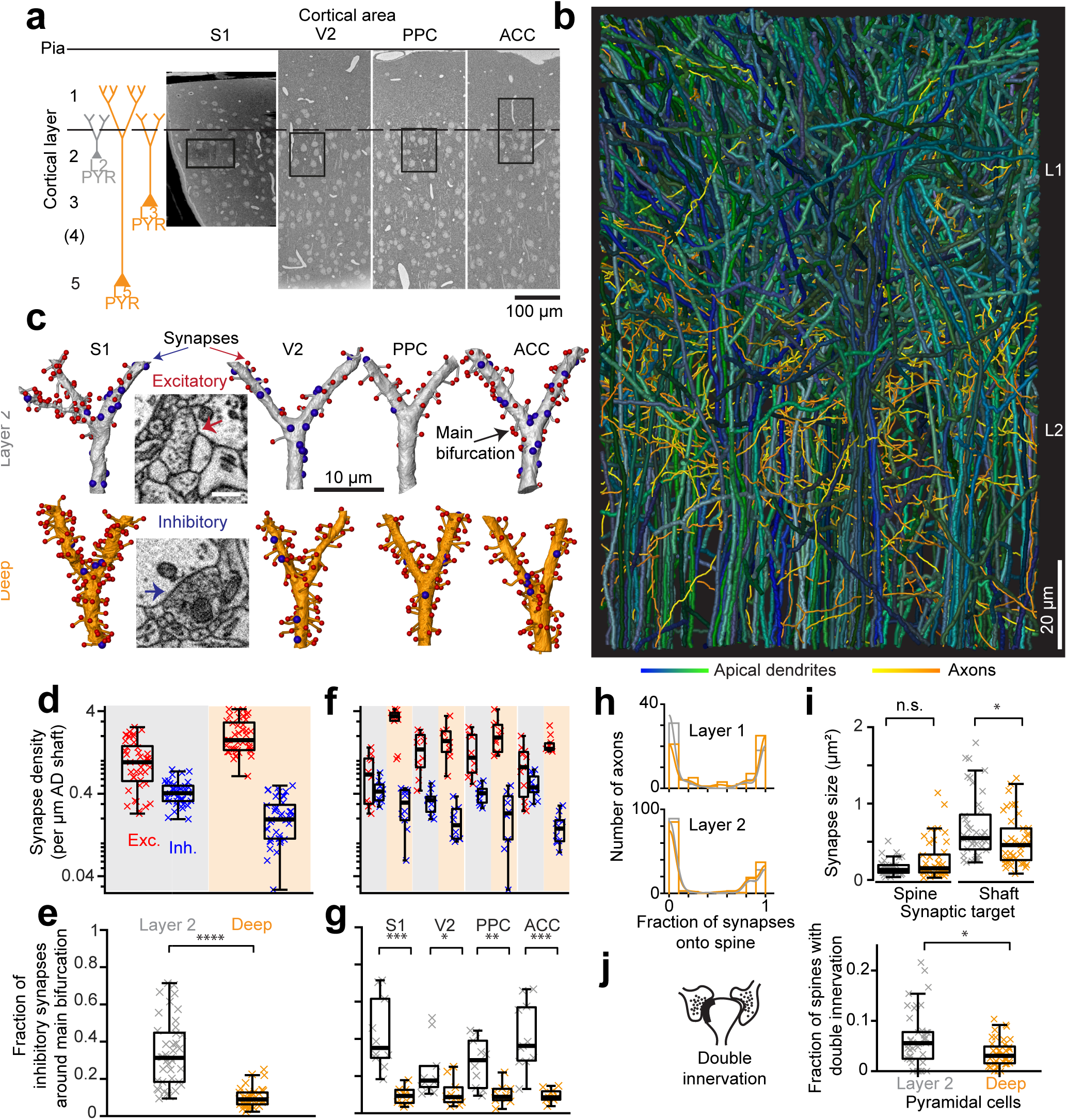
Complete synaptic input mapping of pyramidal cell apical dendrites (ADs) around their main bifurcation. **(a)** Overview EM images indicating the location of 3D EM datasets in primary somatosensory (S1), secondary visual (V2), posterior parietal (PPC) and anterior cingulate (ACC) cortices relative to pia surface (solid line) and layer 1/2 border (dashed line). Schematic location of layer 2 (grey) and deep layer (orange) pyramidal neurons. **(b)** Reconstruction of all ADs contained in the ACC dataset (blue-green, n=61 layer 2 and n=152 deep layer ADs, respectively) and a subset of axons innervating them (n=62, yellow-orange). Note that 80-90% of all pyramidal cells in cortex extend their apical dendrites into the L1/2 border region, allowing for massive synaptic convergence. **(c)** Complete synaptic input maps of apical dendrite main bifurcations for deep layer (orange) and layer 2 (grey) pyramidal cells (example excitatory (spine, red spheres) and inhibitory (shaft, blue spheres) synapses, inset). **(d)** Box plot of inhibitory (blue crosses) and excitatory (red crosses) synapse densities per µm of AD shaft path length for L2 (n=41, left) and DL ADs (n=41, right) in S1, V2, PPC and ACC datasets. Wilcoxon rank-sum test, p<10^-6^, 10^-7^ for excitatory and inhibitory densities, respectively. **(e)** Box plot of fraction of inhibitory synapses at the main bifurcation of deep (orange) and layer 2 (grey) ADs; individual ADs shown (crosses). **(f**,**g)** Same as in **(d**,**e)** separately reported by cortical region (n=20 for S1, V2 and PPC, n=22 for ACC). **(h)** Fraction of synapses made onto spines for axons seeded from synapses on deep (orange) and layer 2 (grey) ADs. Annotations are from layer 1 (n=132, LPtA dataset) and layer 2 (n=289, V2, PPC and ACC datasets); probability density estimations (lines). Note highly bimodal distribution allowing the clear distinction of axons into excitatory (spine-preferring) and inhibitory (shaft-preferring) in upper layers of cortex. **(i)** Size of synapses onto deep (n=41, orange crosses) and layer 2 (n=41, grey crosses) ADs. **(j)** Fraction of double-innervated spines in the main bifurcation annotations (orange, n=41) and layer 2 (grey, n=41). Asterisks indicate significance level of the Wilcoxon rank-sum test (*p<0.05, **p<0.01, ***p<10^-3^, ****p<10^-4^). Scale bars: 0.5 µm (inset in c).

We then identified all the synapses onto 82 ADs within 10 µm around their main bifurcation (Fig. 1c, Suppl. Fig. 1d, n=6240 synapses total, of these n=1092 inhibitory synapses, total dendritic length analyzed: 3.33 mm). To our surprise, the number of inhibitory synapses on DL ADs was exceedingly low (0.22±0.019 inhibitory synapses per µm dendritic shaft path length, mean±SEM, n=41, Fig. 1d,f). Together with the strong excitatory input to these ADs (Fig. 1d,f, 2.12±0.149 synapses per µm dendritic shaft path length), this meant only 9.9±5.1 % (Fig. 1e,g, mean±SD, n=41) inhibitory synapses at the main bifurcations of DL ADs in S1, V2, PPC and ACC cortex. Thus, we did not find an indication of a hot spot of inhibitory inputs at the main bifurcation of DL pyramidal cells that would allow counteracting the regenerative electrical activity observed in apical dendrites (Markram and Sakmann 1994, Markram, Helm et al. 1995, Larkum, Zhu et al. 1999, Major, Larkum et al. 2013).

Next, we compared to ADs from L2 pyramidal cells (Fig. 1c-g). Here, unlike for DL ADs, we found substantial inhibitory input to the apical dendrite at the main bifurcation (Fig. 1d,f, 0.42±0.022 inhibitory synapses per µm dendritic shaft path length, mean±SEM, n=41), with a fraction of 33.6±17.9 % (Fig. 1e,g, mean±SD, n=41) of inhibitory synapses for S1, V2, PPC and ACC. Thus, L2 ADs receive about 3-fold more relative inhibition at their main bifurcation compared to L3 and L5 pyramidal cells (9.9% vs 33.6%, Wilcoxon rank-sum test, p<10^-11^). As controls, we asked whether this difference in inhibitory innervation could be confounded by the way we identified inhibitory vs excitatory synapses: we used the previously reported preferential innervation of dendritic shafts vs spines as criterion for axon identity (Kubota, Karube et al. 2016, Motta, Berning et al. 2018). We found that, in fact, the preference of axons to either innervate shafts or spines of ADs was almost binary (Fig. 1h, Suppl. Fig. 1b), strongly supporting the identification of axons with almost exclusive spine innervation as excitatory and those with almost exclusive shaft innervation as inhibitory. As an additional control, we analyzed synapse size and found it to further enhance, not compensate the observed difference in synapse number (Fig. 1i). Finally, differences in single vs. double-spine innervations between the pyramidal cell types could not account for the observed effect (Fig. 1j).

After having found the strong difference in inhibitory input at the main bifurcation between L2 and the other pyramidal cell types, we next wanted to understand whether the presynaptic inhibitory axons show any innervation selectivity regarding the layer origin of the targeted pyramidal cells’ AD (Fig. 2a-c). This was of special interest since the main bifurcations of all pyramidal cells reside in the same spatial region at the L1/2 border in cortex (Fig. 2d, see also Fig. 1b). We therefore reconstructed presynaptic inhibitory axons that had made at least one synapse onto a pyramidal apical dendrite shaft (Fig. 2a, Suppl. Fig. 2a), identified all other output synapses of these axons, and determined whether these were made again onto the same or different types of pyramidal cell ADs. About 20% of synapses formed by these axons were again established onto apical dendrites (Fig. 2b-c,e, n=183, 20.2±1.1%, mean ± SEM). Surprisingly, the AD innervation showed substantial conditional dependence on the type of AD the axon had been seeded from (Fig. 2b,c), rejecting a model of indiscriminate inhibitory AD innervation: Axons seeded at deeper layer ADs made 15.2±1.6% of the remaining output synapses onto deep apical dendrites (i.e. 74.3% of AD synapses), but only 5.3±0.8 % onto L2 ADs (i.e. 25.7% of AD synapses, Fig. 2b-c, n=91, p<10^-8^, Wilcoxon rank-sum test). Conversely, axons seeded at layer 2 apical dendrites did not innervate their deep layer counterparts to a substantial fraction (Fig. 2b-c, 4.4 ± 0.6%, n=92); Rather, they innervated layer 2 ADs (15.6 ± 1.4 %, i.e. 78.1% of their AD synapses). Since the paths of these inhibitory axons along the depth of the cortex were indistinguishable for those axons targeting deep and layer 2 apical dendrites (Fig. 2d, 18.3 vs. 16.9 mm, n=183), this high connectomic specificity far exceeds average random innervation determined by the trajectory of the presynaptic axons.

**Figure 2.**
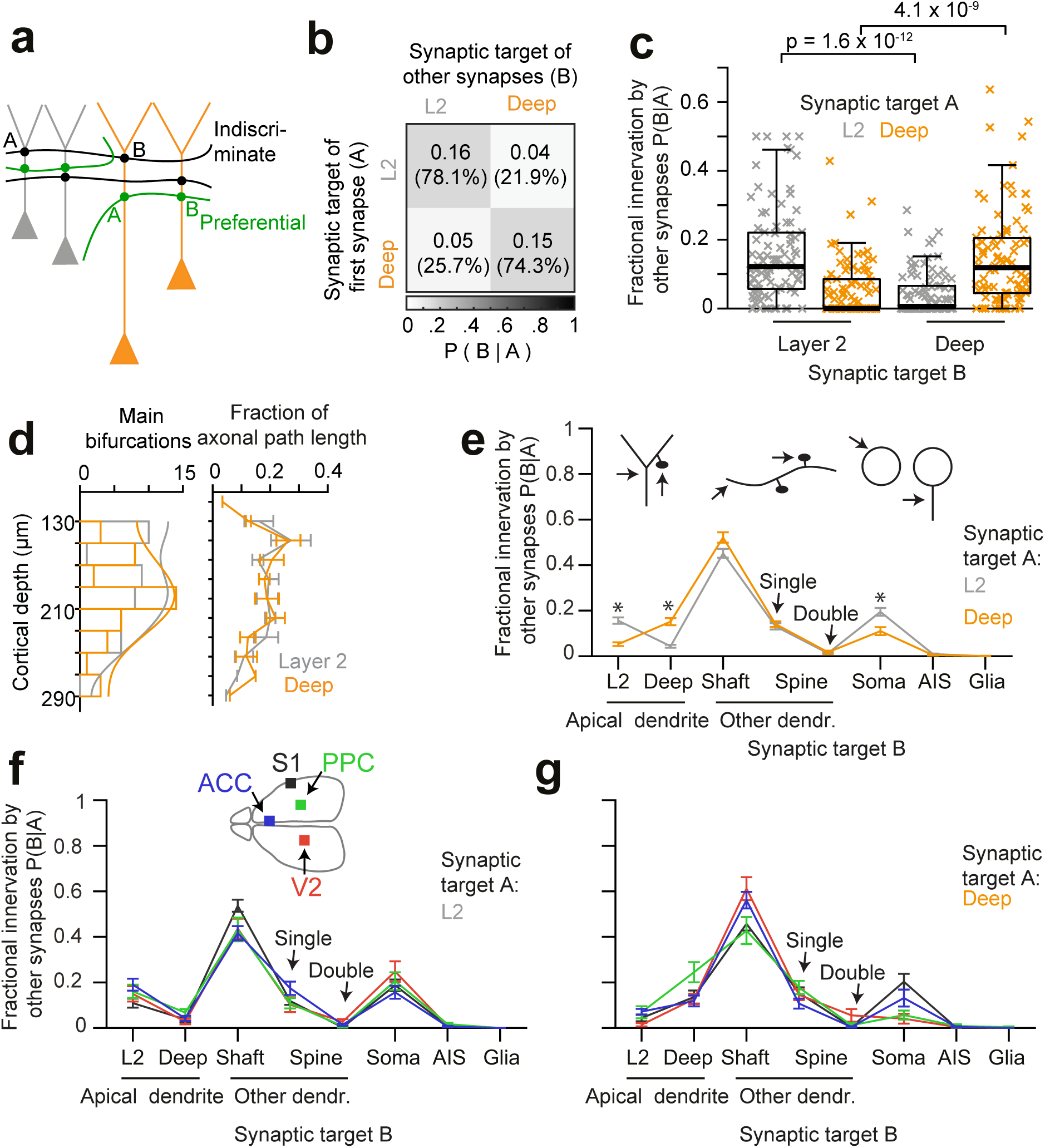
Specificity of inhibitory axons for the type of postsynaptic pyramidal cell. **(a)** Sketch illustrating two extreme innervation models for AD-targeting inhibitory axons: innervation could be selective for the type of AD (L2 vs L3/5 pyramidal cells, green), or rather indiscriminate for the type of AD (black). In the latter case, conditional dependence of targeting p(B|A) would be expected to be absent. **(b)** Conditional dependence of synaptic innervation p(B|A) shown as the mean probability of deep and layer 2 AD targeting (target “B”) given the target of the first synapse of an axon (target “A”). Probabilities are the maximum-likelihood estimate mean of a Dirichlet-multinomial fit to the data. Numbers represent the absolute and fractional (in percent) innervation probability for ADs. **(c)** Box plot of AD fractional innervation for axons seeded from layer 2 (n=92, grey crosses) and deep (n=91, orange crosses) ADs, corresponding to the entries in the innervation matrix (b). P-values are from Wilcoxon rank-sum test. Note that indiscriminate innervation can be revoked. **(d)** Distribution of analyzed main bifurcations along the cortical depth (n=41 per AD type; probability density estimates, lines), and distribution of axonal path length along cortical depth. (n=183 axons, of these n=92 seeded at L2 (gray) and n=91 seeded at DL ADs (orange)). Note that neither the pre-nor the postsynaptic targets are sorted along the cortical axis, excluding simple layering effects for the conditional innervation (b,c). Error bars indicate mean±SEM over cortical region (S1, V2, PPC and ACC). **(e)** Mapping of axonal output onto subcellular targets. Error bars indicate mean±SEM; asterisks: significance of bootstrapping test at p=0.05 with Bonferroni correction. **(f**,**g)** Comparative analysis across cortical regions. Postsynaptic target specificity for axons seeded from (f) layer 2 ADs (n = 21, 20, 21, 30 for S1, V2, PPC and ACC, respectively) and (g) for axons seeded from deep layer ADs (n=19, 20, 20, 32 for S1, V2, PPC and ACC, respectively). Note the high level of quantitative consistency of synaptic target fractions across cortices. Error bars indicate mean±SEM.

We also determined the remaining postsynaptic targets of these AD innervating axons (Fig. 2e), which were remarkably consistent at high quantitative precision over all investigated cortices (S1, V2, PPC and ACC, Fig. 2f,g).

Since we had so far only distinguished layer 2 pyramidal cells from the other pyramidal cell types, we next wanted to determine whether the observed innervation differences were common to L2/3 vs L5 pyramidal cells (thus our data was possibly diluted by L3 pyramidal cells classified as “deep layer”), or whether the inhibitory innervation was in fact distinct between L2 and the other pyramidal cells. We used two datasets in which directly adjacent to the high-resolution EM data volume a low-resolution EM dataset had been acquired that extended down to infragranular layers and allowed the following of ADs to their soma of origin (Fig. 3a, datasets from PPC and LPtA cortex, Suppl. Fig. 1c). The differences in fractional inhibitory innervation were in fact found between L2 pyramidal cells and the other pyramidal cell types, indicating a commonality between L3 and L5 pyramidal cells at their AD main bifurcations (Fig. 3b, inhibitory synapse fraction of 0.38±0.056 vs 0.08±0.008 vs 0.09±0.011 for layers 2, 3 and 5, respectively, mean±SEM, n=30, Kruskal-Wallis test, p<10^-4^).

**Figure 3.**
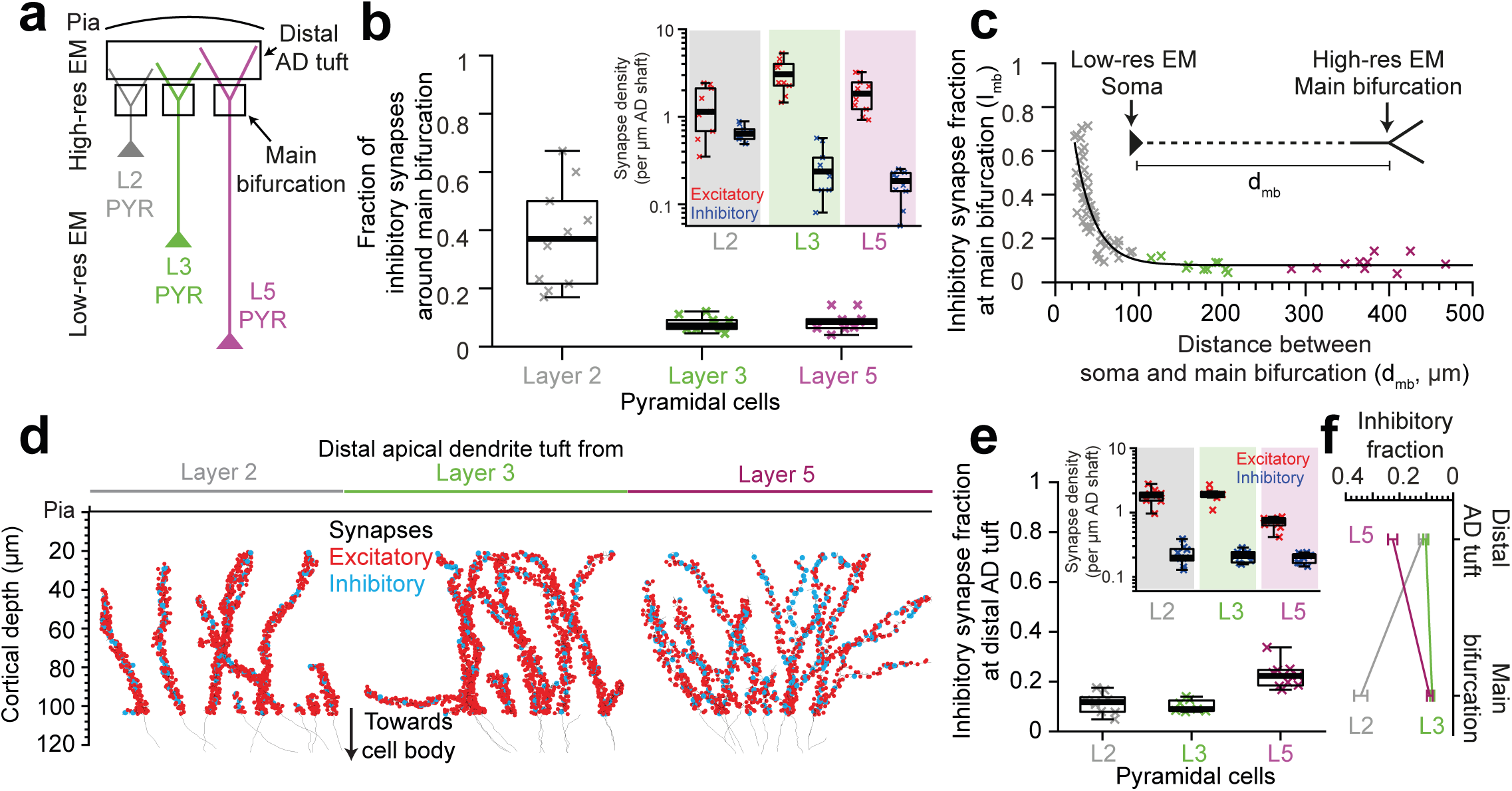
Distinct synaptic input for L2, L3 and L5 pyramidal cells across the upper layers of cortex. **(a)** Synaptic input mapping in EM datasets comprising distal tuft dendrites and main bifurcations, combined with adjacent low-resolution EM datasets to identify soma of origin in L2, L3 or L5 (see Methods). **(b)** Fraction of inhibitory input synapses around the main bifurcation of apical dendrites from layer 2 (grey), 3 (green) and 5 (magenta) pyramidal cells (box plots, n=10 per AD type) and density of excitatory (red crosses) and inhibitory (blue crosses) synapses at the main bifurcation (inset). Note clear distinction of synaptic input composition for L2 vs L3 and L5 pyramidal cells at their main bifurcation. Kruskal-Wallis test, p<10^-4^. **(c)** Relationship between distance of main bifurcation to soma and inhibitory fraction at the main bifurcation for layer 2 (n=51, grey crosses), 3 (n=10, green crosses) and 5 (n=10, magenta crosses) ADs. Black line indicates single exponential regression 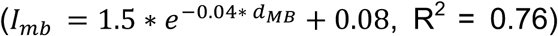. **(d)** Skeleton reconstruction of 11 layer 2, 3 and 5 ADs with all synapses mapped within the high-resolution EM image volume in layer 1 of LPtA cortex (red: excitatory synapses, n=3,782, cyan: inhibitory synapses n=653). Note the difference in synapse densities (red/cyan sphere density) for layer 2, 3, 5 ADs. **(e)** Box plot for fraction of inhibitory synapses at the distal apical dendritic site from layer 2 (n=9 individual branches, grey crosses), 3 (n=7, green crosses) and 5 (n=9, magenta crosses) pyramidal cells and box plot for density of excitatory (red crosses) and inhibitory (blue crosses) synapses at the distal tuft (inset). Note the clear distinction of synaptic input composition in the distal tuft for L2 and L3 vs. L5 pyramidal cells (Kruskal-Wallis test, p<10^-3^). **(f)** Summary of distinct inhibitory input fraction at the main bifurcation and distal tufts of apical dendrites for three main classes of pyramidal cells in the cerebral cortex. Note that only L3 pyramidal cells show a homogeneous balance of inhibitory and excitatory synaptic inputs. Error bars indicate mean±SEM.

We next asked whether the observed cell-type specific inhibitory innervation could be implemented via a synaptic specificity mechanism that was dependent on the dendritic distance of the main bifurcation to the cell body of origin. We found in fact a strong dependence of the fraction of inhibitory synapses on the distance of the main bifurcation to the soma (Fig. 3c, Suppl. Fig. 3a, R^2^=0.76 for single exponential fit, n=71), even within the population of L2 pyramidal cells, suggesting a possible synaptic specificity mechanism that is dependent on the distance to the cell body. Interestingly, this dependence was not found for the absolute density of excitatory or inhibitory synapses, but only for their ratios (Suppl. Fig. 3e, R^2^=0.1, 0.38, respectively).

Finally, we analyzed the inhibitory and excitatory innervation of ADs towards their upper ends in L1 (Fig. 3d,e, LPtA dataset). This was of special interest since the density of dendritic spines in hippocampal pyramidal cells had been previously found to decrease by about 5-fold towards the AD’s tip (Bannister and Larkman 1995, Megías, Emri et al. 2001). We wanted to understand whether this was similar in cortical pyramidal cells, and whether this was accompanied by a drop (or increase) in inhibitory shaft synapse density. We mapped all input synapses onto 11 additional apical tuft dendrites (Fig. 3d, n=4,435 synapses) and found that, first, the excitatory synapse density of L5 pyramidal cells drops 2.75-fold towards their distal tufts. This is not accompanied by a drop in inhibitory inputs, which remains almost constant in density (Fig. 3b,e, Suppl. Fig. 3a-b). Thus, L5 pyramidal cells receive a pronounced inhibitory input peak at their distal dendrites (Fig. 3f, from 8.7±1.1% inhibitory input fraction at main bifurcation to 22.5±1.7% at the distal tips, mean±SEM, n=19 dendritic segments, p<10^-4^ Wilcoxon rank-sum test).

For L2 pyramidal cells, to the contrary, inhibitory synapse density decreases ∼2-fold towards the distal tuft (Fig. 3b,e, 0.47±0.02 vs 0.23±0.03 inhibitory synapses per μm dendritic path, mean±SEM, n=60, Wilcoxon ranksum test, p<10^-4^) while the excitatory input increases by ∼1.5-fold, yielding an about 3-fold reduced inhibitory input fraction towards their distal tufts (Fig. 3b,e, 34.4±2.5% inhibitory input fraction at main bifurcation vs 11.4±1.4% at distal tufts, Wilcoxon rank-sum test, p<10^-4^; see also Suppl. Fig. 3f). Finally, we found that L3 pyramidal cells show no substantial dependence of their synaptic input fractions on the position along the apical dendrite: they have a rather low inhibitory density around their main bifurcation (Fig. 3f, 7.8±0.8%, mean±SEM, n=10, similar to the L5 cells, Wilcoxon rank-sum test, p=0.42), which only slightly increases towards their distal tufts (10.2±0.9%, mean±SEM, n=7 tuft branches, similar to L2 cells, Wilcoxon rank-sum test, p=0.83). With this, we find that the pyramidal cells from layers 2, 3 and 5 show each distinct inhibitory innervation profiles at their distal apical dendrites, providing succinct possibilities for inhibitory control of excitatory inputs in the cortex (Fig. 3f, Suppl. Fig. 3a-c).

## DISCUSSION

We obtained a quantitative connectomic input map of pyramidal cell apical dendrites in several areas of the mouse cerebral cortex. We find distinct profiles of inhibitory to excitatory synaptic innervation balance for all types of pyramidal cells: L2 pyramidal cells receive strong inhibition at their apical dendrites’ main bifurcation, which drops about 3-fold towards the tips of the apical dendrites; in layer 5 pyramidal cells, this profile is inverted with an unexpectedly low fraction of inhibitory inputs at the main bifurcation. Finally, L3 pyramidal cells have a homogeneously low fraction of inhibitory inputs across upper cortex. Furthermore, inhibitory innervation shows high connectomic specificity with axons preferring the innervation of either L2 or deeper-layer pyramidal cells. This suggests a unique innervation profile for each pyramidal cell type, and points to cell-type specific balance of inhibition and excitation in the cortex. These findings were remarkably consistent for all investigated cortices (S1, V2, PPC and ACC).

In physiological experiments, evidence for a strong effect of GABA on regenerative activity in apical dendrites of L5B cells via non-synaptic GABA-B receptors had been reported (Pérez-Garci, Gassmann et al. 2006, Oláh, Füle et al. 2009, Pérez-Garci, Larkum et al. 2013, Abs, Poorthuis et al. 2018). Together with our evidence on low rates of synaptic GABA-ergic innervation at the main bifurcation of these pyramidal cells, this may indicate that L5 pyramidal cells are controlled by overall and slow inhibition at their main bifurcations, but synaptic and specific innervation at their distal dendrites, while L2 pyramidal cells receive specific and synaptic inhibition at their main AD bifurcations. Whether L3 pyramidal cells share the electrical properties of L5 pyramidal cells is still controversial (de Kock and Sakmann 2009, Barth and Poulet 2012). Their synaptic input composition at the main bifurcation as shown here would indicate commonality; but this could be counteracted by different distribution of active transmembrane conductances (Waters, Larkum et al. 2003, Ledergerber and Larkum 2012). Their distinct inhibitory input profile towards the tips of the apical dendrites may suggest that they allow for specific computations not shared by L2 or L5 pyramidal cells.

Spatial distribution of excitatory and inhibitory synapses on the dendritic surface has substantial effects for the integrative properties and output of neurons (Rall 1959, Polsky, Mel et al. 2004, Katz, Menon et al. 2009). Previous studies in hippocampal pyramidal cells had reported a substantial increase in inhibitory synapse fraction towards the distal tuft dendrites which is driven mainly by a large drop in excitatory synapse density (Bannister and Larkman 1995, Megías, Emri et al. 2001, Bloss, Cembrowski et al. 2016). We find that L5 pyramidal cells in cortex exhibit a similar distal inhibitory input domain. Theoretical work indicated that this is potentially a very powerful inhibitory effect (Gidon and Segev 2012). This distal inhibitory domain, however, is absent in the distal tufts of L2 and L3 pyramidal neurons.

Importantly, our data shows a striking specialization of L2 vs L3 pyramidal cells, which has so far evaded analysis (for example, (Iascone, Li et al. 2018)). Only L3, but not L2 or L5 pyramidal cells, keep a roughly even inhibitory to excitatory synaptic input ratio. Recent work using intracellular Gephyrin tagging for the detection of inhibitory synapses and counting of spines for excitatory synapses in LM (Iascone, Li et al. 2018) underestimated the rate of spines by about ∼20% compared to our complete 3D EM based sampling, resulting in an overestimation of the inhibitory-to-excitatory input fraction for L3 pyramidal cells (see also (Chen, Villa et al. 2012)). Our results indicate the importance of separate analysis of L2, L3 and L5 pyramidal cells in cortex (Figs. 1, 3).

In conclusion, our findings describe non-random connectivity at the apical tuft input domain of pyramidal neurons in cortex. They imply unique innervation patterns for the apical dendrites of L2 vs L3 vs L5 pyramidal cells yielding highly variable and cell-type specific inhibitory-to-excitatory synaptic balance, and suggest differential modes of inhibitory operation for the main classes of pyramidal cells in the cortex.

## Author contributions

Conceived, initiated and supervised the study: MH. Performed experiments: AK, JO, FD, KMB; Performed analyses: AK, JO. Wrote the manuscript: AK, MH with contributions by all authors.

## Acknowledgements

We thank Matthew Larkum, Johannes Letzkus and Idan Segev for discussions, Benedikt Staffler, Emmanuel Klinger, Manuel Berning, Alessandro Motta for computational and Jakob Straehle, Meike Schurr, Yunfeng Hua for experimental advice, and Alessandro Motta and Benedikt Staffler for contributing code. We thank M. S. E. A. Aly, L. Bezzenberger, A. B. Brandt, B. Heftrich, A. C. Rix, B. L. Stiehl, C. Arras, C. M. Schumm, D. E. Celik, D. J. Goffitzer, J. Buß, K. M. Trares, K. Weber, L. Buxmann, L. Decker, L. C. R. Kreppner, M. S. Kronawitter, N. M. Böffinger, N. Plath, S. M. Bohne, S. Reichel, T. Engelmann, T. Ernst, T. Winkelmeier, V. C. Kalbert, K. Kramer, L. Präve, M. Präve, N. Berghaus, O. J. Brandt, S. S. Wehrheim for neurite reconstructions, Heiko Wissler, Susanne Babl, Lisa Bezzenberger, Alexander Brandt, Raphael Jakoby, Raphael Kneißl and Marc Kronawitter for annotator training and task management and Heiko Wissler for support with visualizations.

## MATERIALS AND METHODS

### Animal experiments

All experimental procedures were performed according to the law of animal experimentation issued by the German Federal Government under the supervision of local ethics committees and according to the guidelines of the Max Planck Society. The experimental procedures were approved by Regierungspräsidium Darmstadt, under protocol ID V54 - 19c20/15 F126/1015 (LPtA, PPC2) or V54 – 19 c 20/15 – F126/1002 (V2, PPC, ACC). The S1 sample was prepared following experimental procedures approved by Regierung von Oberbayern, 55.2-1-54-2532.3-103-12.

### S1, LPtA sample preparation

The cortical tissue processing, 3D electron microscopy and data alignment for S1 and LPtA samples were described previously in (Berning, Boergens et al. 2015) and (Drawitsch, Karimi et al. 2018), respectively. V2, PPC, ACC and PPC2 samples were processed as follows.

### Transcardial perfusion

Adult (P56 – 57) wild type mice (C57BL/6J) were injected with general analgesia (0.1 mg/kg buprenorphine (Buprenovet, Recipharm, France and 100 mg/kg Metamizol (Metamizol WDT, WDT, Germany)). Next, animals were anesthetized by inhalation of isoflurane (3 – 3.5% in carbogen) and were perfused transcardially using 15 ml of cacodylate buffer (150 mM, Serva, Heidelberg, Germany, pH = 7.4) followed by 30 ml of fixative solution at a flow rate of 10 ml/min. The fixative solution was 2.5% PFA (Sigma-Aldrich, Germany), 1.25% glutaraldehyde (Serva) and 0.5% CaCl_2_ (Sigma-Aldrich) in 80 mM cacodylate buffer (pH = 7.4). The animal was decapitated and the skull was removed to expose the brain. The head was next submerged in fixative solution overnight at 4 °C.

### Cortical region targeting and tissue extraction

First, a 600 µm coronal slice containing the region of interest was acquired using a vibrating microtome (Microm HM 650V, Thermo Fisher Scientific, USA) guided by a reference atlas (Franklin and Paxinos 2008). Next, we used a biopsy punch (1 mm in diameter, KAI medicals, USA) to extract a cylindrical piece of cortex containing layers 1-3 (Suppl. Fig. 1a,c). This tissue was then incubated for 3 - 4 hours (V2, PPC and ACC) or overnight (PPC2) in cacodylate buffer. The PPC2 sample was also subjected to confocal laser scanning light microscopy as previously described (Drawitsch, Karimi et al. 2018).

### En-bloc sample preparation for 3D electron microscopy

Samples were prepared for serial block-face electron microscopy following a slightly modified en-bloc staining method as described in (Hua, Laserstein et al. 2015). In short, cortical tissue was rinsed in cacodylate buffer for 30 min before any staining material was applied. Next, it was transferred into 2% OsO_4_ (Serva, Germany) in cacodylate buffer for 90 min. The sample was then treated with 2.5% ferrocyanide (Potassium hexacyanoferrate trihydrate, Sigma-Aldrich, Germany) in cacodylate buffer for 90 min and 2% buffered OsO_4_ for 45 min. We then rinsed the tissue in cacodylate and ultrapure water (Biochrom, Germany) for 30 min each. Osmium content of the sample was amplified by treatment with saturated aqueous thiocarbohydrazide (TCH, Sigma-Aldrich, Germany) and 2% aqueous OsO_4_ for 60 and 90 min, respectively. The sample was moved to 2% uranyl acetate solution for overnight incubation at 4 °C. The following day, the tissue, still in uranyl acetate, was warmed to 50 °C for 120 min in an oven (Memmert, Germany). This was followed by incubation in lead aspartate at 50 °C for 120 min. The lead aspartate solution was prepared by dissolving 0.066g lead nitrate (Sigma-Aldrich, Germany) in a 10 ml 0.03 M aspartic acid (Serva, Germany) solution and adjusting the pH to 5.0.

Next, the cortical tissue was dehydrated by incubation in 50%-100% ethanol gradient (Serva, Germany). This was followed by at least three 20-45 min incubation steps in pure acetone (Serva, Germany). The sample was then transferred to a 1:1 mixture of acetone and Spurr’s resin (4.1 g ERL 4221, 0.95 g DER 736 and 5.9 g NSA, 113 µl DMAE, Sigma-Aldrich, Germany) for 3-4 h with slow/no rotation. Tubes were opened to allow for acetone evaporation overnight (V2, PPC and ACC samples). At this stage, the PPC2 sample was transferred to a 3:1 mixture of Spurr’s resin and acetone instead. The following day, the infiltration process continued for two 3-hour (PPC2) or one 6-hour (V2, PPC, ACC) incubation steps in pure Spurr’s resin mixture. The tissue was then transferred to a flat-embedding mold and cured at 70 °C for at least 48 hours. Specific time and temperature of dehydration and embedding steps are detailed in Suppl. Table 2.

Note that all procedures were performed at room temperature in 2 ml reaction tubes (Eppendorf, Germany) unless stated otherwise. In addition, the initial incubation steps (until dehydration) were performed with the aid of an automatic microwave tissue processor (Leica EM AMW, Leica, Germany) for V2, PPC, and ACC samples. Finally, treatment steps were interleaved with two 30 min washing steps in ultrapure water (from TCH step until dehydration initiation).

### Serial block-face electron microscopy (SBEM)

The samples were excised from the resin block and mounted on an aluminum pin using epoxy glue (Uhu plus schnellfest, Uhu, Germany). They were then trimmed to a block-face area of ∼ 750 µm × 750 µm using a diamond head trimming machine (Leica EM TRIM2, Leica, Germany). In addition, tissue was sputter coated with 100 - 200 nm of gold (Leica ACE600 Sputter Coater, Leica, Germany) to increase conductivity and reduce charging artefacts.

The SBEM microtome (courtesy of W. Denk) was fit inside the door of a scanning electron microscope (Suppl. Table 1, FEI, Thermo Fisher Scientific, USA). The microtome and microscope were controlled using custom written software during the volume imaging process; focus and stigmation were corrected manually (V2, PPC, ACC and LPtA) or using custom written auto-correction routines (S1, PPC2). The region of interest was imaged using overlapping image tiles and cutting direction was along the tangential (S1, V2, PPC, ACC, PPC2) or radial (LPtA) axes of cortex. The region of interest was targeted to an area close to layer 1/2 border (S1, V2, PPC and ACC, Fig. 1a). The final imaged volume and the nominal voxel size is detailed in Suppl. Table 1. Note that the LPtA and PPC2 datasets were adjacent to datasets extending to the middle of layer 5 imaged at lower resolution (22.48×22.48×30 and 44.96×44.96×120 nm^3^ for PPC2 and LPTA, respectively).

### Image Alignment

The alignment for PPC, V2 and ACC dataset was done using custom-written MATLAB (Mathworks, USA) routines. We used cross-correlation or speeded-up robust feature (SURF) detection in the overlap region to measure the relative shift between image patches. These patches were full image tiles (V2, ACC) or tile sub regions (PPC, 256 × 256 pixels). The position of each patch was then globally optimized using least-square regression. The image volume was then partitioned into 1024×1024×1024 voxel blocks and written into webKnossos (Boergens, Berning et al. 2017) three-dimensional format. The dataset was then transferred to the data store accessible by webKnossos for in-browser neurite skeleton reconstruction and synapse annotation.

The PPC2 dataset was aligned using the affine alignment method from (Scheffer, Karsh et al. 2013). The routines were modified to give sub regions of image tiles unique affine transformations. In addition, methods to exclude featureless blood vessel and nuclei were improved.

### Reconstruction and synapse annotation analysis

The skeleton reconstruction and synapse annotations were saved as a NML (XML-based format) file. They were subsequently parsed into a MATLAB (release 2017b) class with node and edge list properties using custom-written C++ routines. Each node had accompanying attributes, such as, comment string and coordinate. These attributes were used to extract different features of the annotation as described in the following sections.

### Apical dendrite (AD) definition and classification

Apical dendrites (ADs) were identified based on their radial direction and diameter (∼1-3 µm). The soma morphology and axon initial segment direction (towards white matter (WM)) of candidate pyramidal cells were examined where possible.

Within the S1, V2, PPC and ACC datasets, ADs were classified depending on the existence of soma in the image volume (Fig. 1a). ADs with soma in the image volume were classified as layer 2 (L2) and other apical dendrite were classified as deep layer (L3/5) ADs.

The depth of the apical dendrite’s source soma relative to pial surface was used to differentiate layer 2, 3 and 5 cells in the LPtA and PPC2 datasets which contained all these layers. Layer 5 (L5) apical dendrites were from neurons at a depth of at least 620 µm and 500 µm from pia for LPtA and PPC2, respectively. Layer 2 apical dendrites (L2) had a soma depth of 190 - 270 µm and 200 - 300 µm in LPtA and PPC2, respectively. Finally, layer 3 (L3) apical dendrites arise from the cortical depth between L2 and L5. Note that, LPtA and PPC cortical regions do not possess a prominent layer 4 (Kolb and Walkey 1987).

The difference between nominal and actual cutting thickness in the LPtA dataset was resulting in apparently ellipsoid somata (compressed along the cutting axis). To obtain an accurate estimate of soma depth relative to pial surface, the section thickness was corrected (by a factor of 1.49), assuming soma dimensions to be similar in-plane and along the cutting direction.

The main bifurcation of apical dendrites was defined as the branching point with two daughter branches of similar thickness and branching angle (resulting in a “Y” shape, Fig. 1c).

We used the ACC dataset to reconstruct all ADs (Fig. 1b, blue-green). They were detected by examining the border of dataset facing WM for deeper layer ADs (n=152) and by identifying all L2 pyramidal neurons contained for L2 ADs (n=61). The AD locations were used as seed points for manual annotation ignoring spines.

### Complete synaptic input mapping of apical dendrites

Apical dendrites and their associated spines were skeleton reconstructed and the synapses on their shaft and spines were annotated either within a bounding box of size 20 × 20 × 20 µm^3^ (Fig. 1c-g, 3b-c, Suppl. Fig. 1d, 3a, n = 20 for S1, V2, PPC, n=30 for PPC2, n=22 for ACC) around the main bifurcation or throughout the dataset (Fig. 3d-e, Suppl. Fig. 3a, n=11 for LPtA, n=6 for ACC, n=4 for V2, PPC and ACC) by a neuroscientist expert annotator (JO or AK). Synapses were identified within the SBEM data based on the presence of vesicle cloud and postsynaptic density as described previously (Schmidt, Gour et al. 2017). We also annotated spine neck synapses and double innervations of spine heads by two axonal boutons (Fig. 1j, (Kubota, Hatada et al. 2007)). Shaft synapses, synapses on spine necks and secondary spine innervations were treated as inhibitory synapses. Primary spine innervations were treated as excitatory synapses (Fig. 1,3). All synapse annotations were validated by an additional expert annotator, and only synapses where annotators agreed were used for analysis.

The fraction of inhibitory synapses was defined as the number of inhibitory synapses divided by the total number of synapses (Fig. 1e,g, Fig. 3b,e). The inhibitory and excitatory synapse density was calculated by dividing the number of synapses by the path length of the apical dendrite shaft (Fig. 1d,f, insets in Fig. 3b,e). Path length was measured by removing the spine necks from the skeleton reconstruction and summing the lengths of the remaining edges. Note that inhibitory input was measured separately for each distal tuft branch in the LPtA dataset resulting in 9, 7 and 9 dendritic segments for L2, 3 and 5 cells, respectively (Fig. 3d-f).

### Inhibitory input fraction mapping in upper cortex

We first estimated the approximate location of each dataset relative to pia (Fig. 1a, 125, 215, 170, 110, 10, 20 µm for S1, V2, PPC, ACC, PPC2 and LPtA, respectively) based on their position in coronal overview images and transformed all reconstructions into a common coordinate system (Suppl. Fig. 3a). Next, we partitioned these reconstructions (n=131, total AD shaft path length=13.5 mm) into virtual 100 µm thick cortical tangential sections. This process resulted in dendritic segments at each depth bin. We then combined synapse counts and path lengths from all segments within each virtual cortical section to calculate the gross average of inhibitory fraction and synapse densities (lines in Suppl. Fig. 3b-c). We also used the 95% bootstrap confidence interval to estimate the dependence of the gross average on the sample composition (shades in Suppl. Fig. 3b-c, n=10,000 bootstrap samples).

### Soma, main bifurcation distance effect on synapse composition at the main bifurcation

We annotated the apical trunk connecting the soma to main bifurcation within the high- and low-res EM data volumes for the L2 (n=51, S1, V2, PPC, ACC, PPC2), L3 (n=10, PPC2) and L5 (n=10, PPC2) pyramidal cells and considered the path length of these annotations as the soma to main bifurcation distance (Fig. 3c, Suppl. Fig. 3e).

### Inhibitory fraction along the AD of L2 pyramidal neurons

We binned the excitatory and inhibitory synapses based on their path distance to soma in the L2 pyramidal ADs (Suppl. Fig. 3f, n=59, n=12,395 synapses, bin size=10µm). This allowed us to measure the fraction of inhibitory synapses as a function of dendritic path distance to the center of soma (10-330 µm distance range). Soma distance bins with less than 4 synapses were merged to their immediate neighbor that contained at least 4 synapses. This was to avoid extreme values introduced by computing the inhibitory ratio in a bin with low synapse numbers.

### Spine innervation fraction distribution

We randomly selected axons targeting a shaft or a spine of ADs and annotated at least 6 other synapses they formed within the image volume. We then determined whether the other postsynaptic targets are primary spine innervations (this excluded shaft, spine neck and secondary spine innervation). Finally, we plotted the histogram (bin size=0.1) and the probability density estimate (bandwidth = 0.0573) of spine innervation fraction for each synapse type (Suppl. Fig. 1b) or combined the data based on the seed AD type (Fig. 1h, n = 132 and 289 for datasets in layers 1 and 2, respectively).

### Synapse size estimation

A random subset of shaft and spine (n = 41 per AD type) synapses were used to determine the synaptic interface area (Fig. 1i, S1, PPC, V2 and ACC). For this, an expert annotator placed two edges along the longest dimension of the synapse and its approximate orthogonal direction. These two edges were used as minor and major axes of an ellipse to estimate the contact area (*area* = *π* * *semi* − *major axis* * *semi* − *minor axis*).

### Distribution of main bifurcations and axonal paths along upper cortex

We obtained the main bifurcation depth relative to pia by transforming the datasets into a common coordinate system as described above (n = 82, Fig. 1c, main bifurcation annotations). Next, we used this to create a histogram (bin size of 20 µm) and probability density estimate (bandwidth: 25 µm) along the cortical depth (Fig. 2b, left panel) for main bifurcation densities. Note that this is only a subset of main bifurcations within each image volume (See Fig. 1b).

We used these datasets to also obtain a density estimate for inhibitory axonal paths. For this, we removed the nodes used for marking synapse locations in the inhibitory axonal reconstructions and computed the average fraction (across S1, V2, ACC and PPC datasets) of the remaining axonal path within 20 µm tangential cortical slices (Fig. 2b, right panel).

### Fractional innervation of inhibitory axons

We selected a random subset of inhibitory synapses from layer 2 (n = 21, 20, 21, 30 for S1, V2, PPC and ACC) and deep layer (n = 19, 20, 20, 32 for S1, V2, PPC and ACC) ADs. An expert annotator (JO or AK) then reconstructed the axons throughout the dataset. We also annotated all the other synapses and postsynaptic partners for each axon. In addition, reconstructions were checked by another expert annotator (JO or AK) for morphological irregularities such as sharp branching angles and untraced endings.

The postsynaptic targets were categorized into one of the following: layer 2 or deep layer apical dendrite (shaft and spine of trunk and primary branches), shaft of other dendrites, single- or double-innervated (including neck targeting) spine, layer 2 cell body, axon initial segment (AIS) or glia. The innervation fraction of an axon was computed by dividing the number of synapses for each specific target by the total number of synapses of that axon (the seed synapse was excluded). The innervation fractions were averaged for each AD seed type (Fig. 2e). Each dataset average was also computed separately (Fig. 2f - g).

### Dirichlet-multinomial model for postsynaptic targeting probability

A Dirichlet-multinomial model was chosen as the generative process for the axonal postsynaptic target counts. In this model, a multinomial probability vector is drawn from a Dirichlet distribution for each axon. Synapse targeting count is then a multinomial sample of this probability vector. We then used a Newton-iteration method (Minka 2000) to find the maximum-likelihood estimate of the Dirichlet-multinomial distribution of our axonal targeting counts. We then used the mean of this distribution as the probability of axons seeded from layer 2 and deep ADs to target each AD type. In addition, we reported the fraction (in percent) of AD targeting probability (Fig. 2b).

### Visualizations of dendritic surfaces

To visualize the surface of the main bifurcation of PPC and S1 apical dendrites and axons, we segmented the image volume using SegEM (Berning, Boergens et al. 2015) and collected all the segments compromising the dendritic shafts or axons. The volume data was then imported to MATLAB, binarized and smoothed using a (9 × 9 × 9) voxel Gaussian convolution kernel with standard deviation of 8 voxels. The isosurface was then constructed at a threshold of 0.2 (Fig. 1c, Suppl. Fig. 1d, Suppl. Fig. 2a).

Volume of the AD shaft at main bifurcation in V2 and ACC datasets were generated by tracing the outline of the dendrites using the volume tracing mode in webKnossos and 3D data was processed in a similar fashion.

In Fig. 1c, the surface of each main bifurcation was overlaid with skeleton reconstruction of spine necks and spheres demonstrating synapses. The visualizations were generated using Amira (Thermo Fisher Scientific, USA).

### Statistics

Significance of difference between L2 and DL AD excitatory and inhibitory synapse densities (Fig. 1d,f), inhibitory fractions (Fig. 1e,g), synapse size (Fig. 1i), double innervation of spines (Fig. 1j) and AD innervation fraction (Fig. 2c) were tested using the non-parametric Wilcoxon rank-sum test. In addition, the rank-sum test was used to test for the difference of inhibitory fraction and excitatory/inhibitory synapse densities between the main bifurcation and the distal tuft area for each pyramidal cell type (Fig. 3b,e-f). Finally, the difference in the inhibitory fraction between L2, 3 and 5 apical dendrites (n=3 independent samples) at the main bifurcation and on the distal tufts were tested using the non-parametric Kruskal-Wallis test to account for the small sample sizes (n=10, 10, 10 main bifurcations and n=9, 7, 9 distal tuft branches for L2, 3 and 5, respectively. Fig. 3b,e).

To identify significant differences in the axonal innervation fractions, a bootstrapping test was used. The specificities of the two axonal groups were concatenated and 10,000 bootstrap resamples were drawn to match the number of axons seeded from L2 and DL ADs. The mean difference in each bootstrap resample between L2 and DL groups was compared to the sample mean difference. The p-value was defined as the fraction of bootstrap resamples which had a value more extreme compared to the sample mean. The significance threshold was set to 0.05 with Bonferroni correction for 8 comparisons (Fig. 2e).

To investigate the relationship between synaptic composition at the main bifurcation and distance to soma, single-term exponential models (with offset) of the form *y* = *c* + *a* * *e*^*bx*^ were fit to the data from layers 2-5. The coefficient of determination (R^2^) was computed as a goodness of fit measure.

### Data and software availability

All 6 datasets will be made available upon publication. All software used for analysis will be made public under the MIT license:

