## Supplementary material for "Cell-type specific innervation of cortical pyramidal cells at their apical tufts"

### SUPPLEMENTARY TABLES

#### Supplementary table 1

| | S1 | V2* | PPC* | ACC | LPtA | PPC2 |
|---|---|---|---|---|---|---|
| **Mouse age (postnatal days)** | 28 | 56 | 56 | 56 | 46 | 57 |
| **Sample location (in mm) relative to bregma (AP,ML)** | (-1.2, -3.9) | (-2.6,1.7) | (-2,-1.7) | (0.8,-0.15) | (-2.1, -1.3) | (-2, -1.5) |
| **Sample section thickness (µm)** | 1000 | 600 | 600 | 600 | 1000 | 600 |
| **SEM type (FEI, USA)** | Magellan | Quanta | Quanta | Quanta | Verios | Magellan |
| **Electron landing energy (KeV)** | 2.8 | 2.8 | 2.8 | 2.8 | 2.8 | 2.8 |
| **Beam current (nA)** | 3.2 | 0.2 | 0.2 | 0.11-0.2 | 0.8 | 1.6 |
| **Beam dwell time (µs)** | 0.1 | 2.1 - 2.8 | 2.3 | 2.1 - 2.8 | 0.5 | 0.2 |
| **Tile configuration in plane (x, y)** | 3 x 3 | 1 x 2 | 1 x 2 | 1 x 3 | 4 x 5 | 5x5, 5x10 |
| **Overlap between tiles (x, y)** | (21%, 11%) | (N/A, 8%) | (N/A, 6%) | (N/A, 4.5%) | (8%, 12%) | (6%, 7%) |
| **Single tile resolution (pixels)** | 3072x2048 | 6144x4096 | 6144x4096 | 6144x4096 | 3072x2048 | 4096x3536 |
| **voxel size ($nm^3$)** | 11.24x11.24x28 | 12x12x30 | 12x12x30 | 12x12x30 | 11.24x11.24x30 | 11.24x11.24x30 |
| **Final high-resolution volume ($\mu m^3$)** | 66x89x202 | 72x91x153 | 72x93x141 | 70x141x98 | 130x110x85 | 200x(185-370)x200 |
| **Dataset distance from pia (µm)** | 125 | 215 | 170 | 110 | 20 | 10 |
| **Low-resolution EM volume existence**** | | | | | X | X |
| **Staining approach** | Manual | AMW assistance | AMW assistance | AMW assistance | AMW assistance | Manual |
| **Staining protocol** | Conventio-nal | Modified Hua | Modified Hua | Modified Hua | Modified Hua | Modified Hua |

* From opposing hemispheres of same animal

** Used for finding the apical dendrite's cell body of origin

**Experimental details.** This table summarizes experimental parameters used for sample preparation and volumetric electron microscopy in 6 datasets from 5 cortical regions.

### Supplementary table 2

| | V2 | PPC | ACC | PPC2 |
|---|---|---|---|---|
| **50% Ethanol** | 30 min @ 4°C | 30 min @ 4°C | 30 min @ 4°C | 30 min @ RT (cooled) |
| **75% Ethanol** | 45 min @ 4°C | 45 min @ 4°C | 45 min @ 4°C | 30 min @ RT (cooled) |
| **100% Ethanol** | 45 min @ RT | 45 min @ RT | 45 min @ RT | 2 times, 30 min @ RT |
| **Pure acetone** | 3 times , 45 min each @ RT | 3 times , 45 min each @ RT | 3 times , 45 min each @ RT | 4 times, 20 min each @ RT |
| **50% Spurr's resin in acetone** | 3 hr @ RT (no rotation, closed tube). Next, open tube for 90 min initial rotation + overnight @ RT | 3 hr @ RT (no rotation, closed tube). Next, open tube for 90 min initial rotation + overnight @ RT | 3 hr @ RT (no rotation, closed tube). Next, open tube for 90 min initial rotation + overnight @ RT | 4 hr @ RT, closed tube cap, slow rotation |
| **75% Spurr's resin** | N/A | N/A | N/A | Overnight @ RT. Slow rotation, closed caps. |
| **100% Spurr's resin** | 6 hr @ RT | 6 hr @ RT | 6 hr @ RT | 2 times, 3 hr @ RT each. No rotation |

**Dehydration and embedding times and temperatures**. Time and duration of each dehydration and embedding step for samples from 3 cortical region (n=4).

### Supplementary table 3

| | S1 | V2* | PPC* | ACC | LPtA | PPC2 |
|---|---|---|---|---|---|---|
| **Main bifurcation input mapping (Fig. 1c-g, Fig. 3b-c)** | X | X | X | X | | X |
| **Dense apical dendrite reconstruction (Fig. 1b)** | | | | X | | |
| **Synapse size estimation (Fig. 1i)** | X | X | X | X | | |
| **Spine innervation fraction (Fig. 1h)** | | X | X | X | X | |
| **Double spine fraction (Fig. 1j)** | X | X | X | X | | |
| **Fractional innervation of inhibitory axons (Fig. 2)** | X | X | X | X | | |
| **Cell type comparisons of inhibitory fraction (Fig 3)** | X (only L2) | X(only L2) | X(only L2) | X(only L2) | X | X |
| **Path distance to soma dependency (Fig. 3c, Suppl. Fig. 3e-f)** | X (only L2) | X(only L2) | X(only L2) | X(only L2) | X | X |
| **Profile of inhibitory fraction along upper cortex(Suppl. Fig. 3a-d)** | X | X | X | X | X | X |

**Data analysis**. Summary of analyses carried out in all 6 datasets.

### SUPPLEMENTARY FIGURES

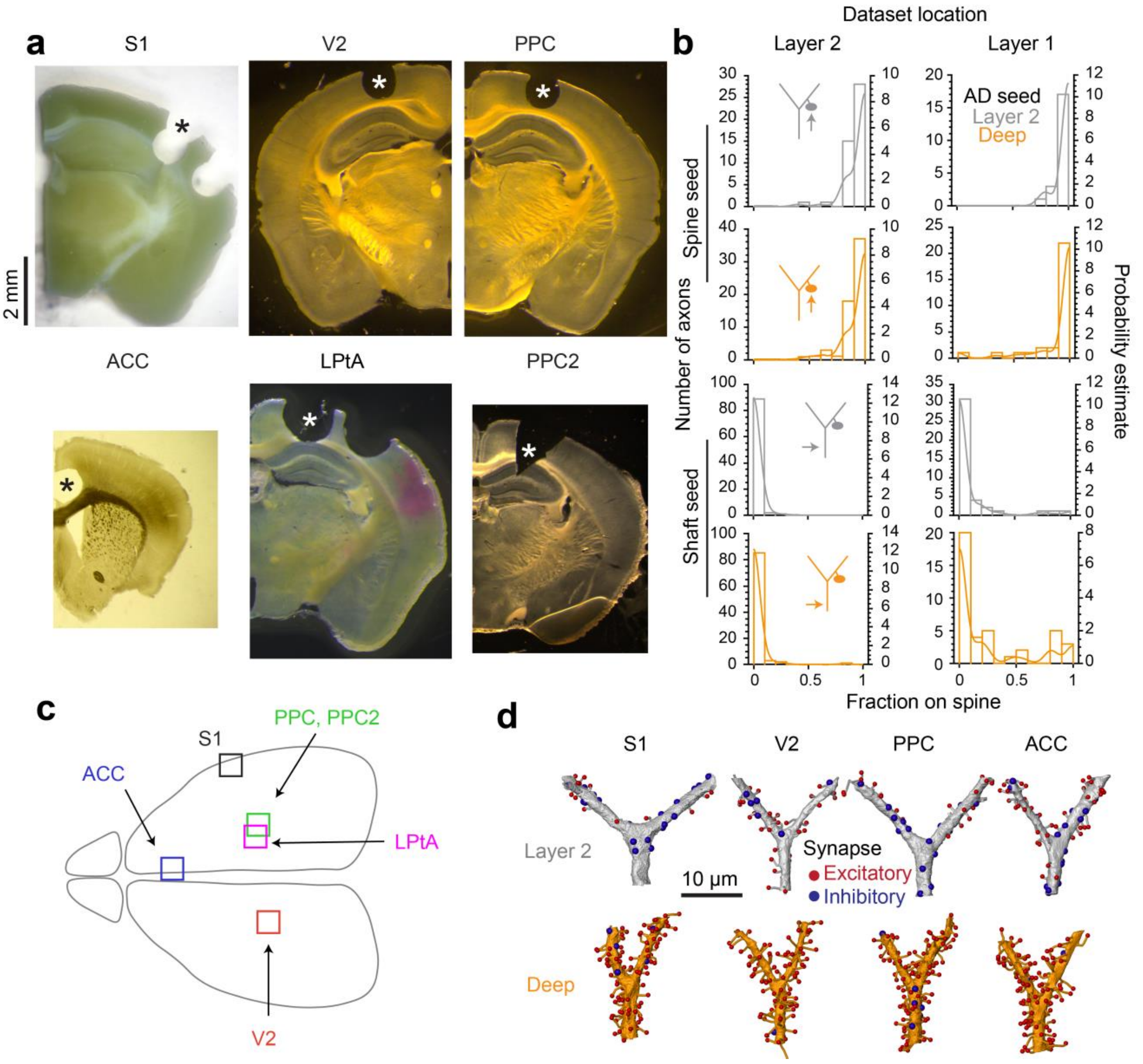


**Supplementary Figure 1**

**Cortical region location, distribution of spine targeting fraction and complete synaptic input maps at main bifurcation. (a)** Coronal sections (thickness: 100-1000 µm) demonstrating the cortical regions (asterisks) used for 3D electron microscopy. **(b)** Histogram and probability density estimates (lines) of primary spine targeting fraction for axons seeded from spine and shaft of ADs in layer 2 (V2, PPC and ACC) and layer 1 (LPtA) cortical regions. See also Fig. 1h. **(c)** Schematic of mouse brain demonstrating approximate location of all 6 cortical regions used for 3D-EM. **(d)** Additional complete synaptic input maps around the main bifurcation. See Fig. 1c.

**a**

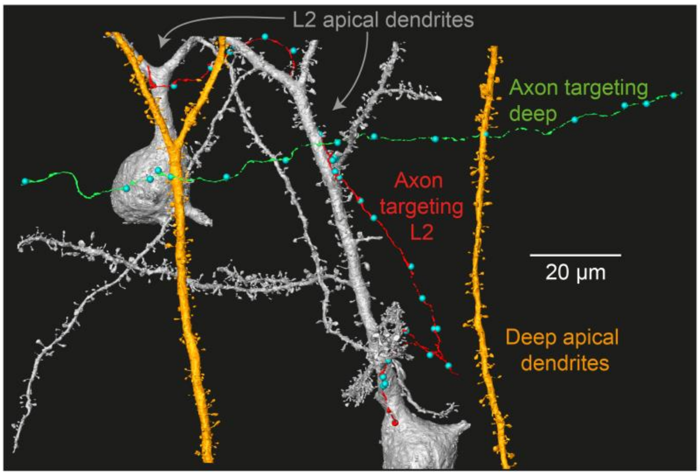


**Supplementary Figure 2**

**Reconstruction of axons and their target layer 2 and deep apical dendrites. (a)** Two example axons (red: targeting layer 2, green: targeting deep) specific for layer 2 (grey) and deep layer (orange) apical dendrites. Blue spheres indicate the location of all output synapses of the axons in the volume.

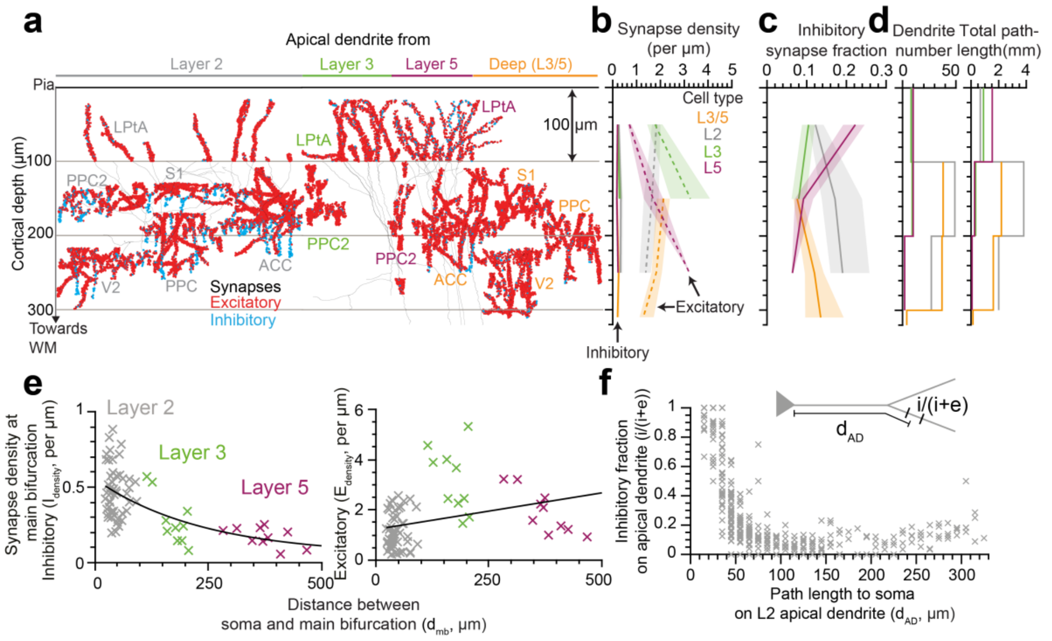


### Supplementary Figure 3

**Synpatic composition for layer 2, 3 and 5 apical dendrites (ADs) along the upper cortex and the relationship between path distance to soma and synapse densities (L2, 3 and 5) at the main bifurcation or fraction of inhibitory synapses along the AD (only L2). (a)** Partial or complete skeleton reconstruction of 131 layer 2, 3 and 5 ADs with all synapses mapped (red: excitatory, n=22,499, cyan: inhibitory n=3,579, total AD shaft path length=13.5 mm). 5 datasets containing layer 2 (n = 20, 20, 22, 28, 30 for S1, V2, PPC, ACC and PPC2, respectively) and a dataset from layer 1 (n=11, LPtA) was used. Note the difference in synapse densities (red/cyan sphere density) for layer 2, 3 , 5 and deep (L3/5) ADs at different cortical depths relative to pia. Horizontal location of the skeleton tracings is adjusted for illustration. **(b)** Spatial distribution of excitatory (dashed line) and inhibitory synapse (solid line) densities of L2 (gray), 3 (green), 5 (magenta) and 3/5 (deep, orange) ADs in upper cortex (bin size=100µm). Lines and shades indicate the gross average and 95% bootstrap confidence interval (n=10,000 resamples) for data combined across all datasets, respectively. **(c)** same as in **(b)** for fraction of inhibitory synapses. **(d)** Histogram of the number of dendrites and total AD shaft path length within each cortical depth bin. **(e)** Relationship between distance to soma and inhibitory and excitatory synapse densities at the main bifurcation for layer 2 (n=51, grey crosses), 3 (n=10, green crosses) and 5 (n=10, magenta crosses) ADs. Black line indicates single exponential fits for inhibitory ($I_{density} = 0.51 * e^{-0.004 * d_{MB}} + 0.048$, $R^2$=0.38) and excitatory ($E_{density} = 1332.6 * e^{2.1 * 10^{-6} * d_{MB}} - 1331.4$, $R^2$=0.1) synapses. **(f)** Relationship between path distance to soma on the apical dendrite and the fraction of inhibitory synapses on L2 apical dendrites (n=59, S1, V2, PPC, ACC, PPC2 and LPtA datasets, n=12,395 synapses). Each cross represents the inhibitory fraction (i/(i+e)) for a single apical dendrite within a 10 μm path length range ($d_{AD}$) to cell body of origin. Also see schematics (inset).
